# Chaotic synchronization in adaptive networks of pulse-coupled oscillators

**DOI:** 10.1101/2024.07.11.603061

**Authors:** Germán Mato, Antonio Politi, Alessandro Torcini

## Abstract

Ensembles of phase-oscillators are known to exhibit a variety of collective regimes. Here, we show that a simple mean-field model involving two heterogenous populations of pulse-coupled oscillators, exhibits, in the strong-coupling limit, a robust irregular macroscopic dynamics. The resulting, strongly synchronized, regime is sustained by a homeostatic mechanism induced by the shape of the phase-response curve combined with adaptive coupling strength, included to account for energy dissipated by the pulse emission. The proposed setup mimicks a neural network composed of excitatory and inhibitory neurons.

---

Understanding the behavior of large ensembles of coupled oscillators is the objective of active research since many years [1, 2]. This includes diverse areas such as engineering, neuroscience, and systems biology [3–6]. One of the key points is the spontaneous emergence of various forms of synchronization [7, 8], i.e. of collective regimes out of given microscopic rules. Identifying the relationship between the microscopic and macroscopic world is, however, non trivial and a general theory is still lacking. In this Letter, we focus on the most challenging phenomenon: collective chaos (CC), a stochastic-like, partially shynchronized regime. The emergence of CC in ensembles of oscillators which behave chaotically is a well established fact both in heterogeneous [9] and homogenous [10–12] networks.

Pulse-coupled phase oscillators can give rise to quite complex scenarios. Low dimensional CC has been reported in two coupled populations of spiking neurons [13], and of Winfree oscillators [14], where it has been interpreted as a sort of chimera mechanism [15]. With reference to a single population, low-dimensional CC was found in quadratic integrate-and-fire (QIF) neurons in [16], where the delay plays a crucial role, by increasing the phase space dimensionality. Finally, high- (possibly infinite-) dimensional CC was reported in [17, 18], where heterogeneity appears to be a crucial ingredient.

In all such cases, the connectivity is proportional (if not equal, as in mean-field models) to the number of oscillators, while the coupling strength is assumed to scale as 1*/N*, to avoid unpleasant divergencies in the thermodynamic limit. We call this setup as S1. However, there exists a second setup (S2), inherited form the study of spin glasses, where the coupling strength is assumed to be of order 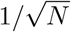 [19, 20]: it makes sense whenever a number of positive and negative coupling terms proportional to *N* tend to cancel (balance) each other, suppressing divergencies. The setup S2 is quite popular within neuroscience since the balance between excitatory and inhibitory synaptic inputs spontaneously emerges in the presence of strong external currents [21], and very recently, it was proven to work also when the currents are weak [22], if synaptic depression is included. Besides, while the microscopic dynamics is almost regular in S1, it is strongly erratic in S2, as expected for the brain neural activity [23].

In this Letter, we revisit the S1 setup, by studying the dynamics of two coupled populations of heterogeneous pulse-coupled phase oscillators (conveniently termed excitatory and inhibitory neurons). An important novelty is the inclusion of an adaptive mechanism – short term depression (STD) – proposed in [24, 25], which limits the firing activity of the neurons taking in account the limitation of the resources. This mechanism gives rise to additional nonlinearities that persist in the infinite size limit without loosing microscopic variability, as shown by the mean field theory developed in [26].

In our model, oscillator interactions are mediated by phase response curves (PRCs), as widely assumed in computational neuroscience [5, 27–29]. Finally, we consider a globally coupled network, finding that above a critical value *G*_*θ*_ of the coupling strength *G*, a fluctuationless asynchronous regime (AR) destabilizes, giving rise to CC accompanied by a highly fluctuating microscopic dynamics. The regime is kept in a balanced state by a self-adjusting synchronization which progressively enhances upon increasing the coupling strength. This is the result of a non trivial synthesis of STD and the PRC shape: an increased level of synchrony among the oscillators leads to a decrease in the effective interaction, because of the vanishing amplitude of the PRC in proximity to the pulse emission.

## Network Model

As shown in [3, 22, 28, 30], it is possible to derive analytically phase oscillator models describing the evolution of the membrane potentials of pulse-coupled supra-threshold neurons, whenever the neuron model is one dimensional. In particular, by following [22] the evolution of the phase 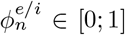 of the *n*-th neuron (*e/i* denotes the excitatory/inhibitory nature) can be written as

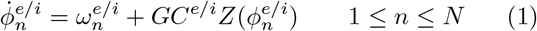

where the two populations are assumed to be characterized by the same number of neurons. Whenever the phase variable 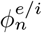 reaches 1, an instantaneous pulse (a *δ*-spike) is delivered to all the excitatory and inhibitory oscillators and the phase is reset to zero. The natural frequencies 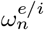 are randomly selected, according to a generic distribution with a finite support 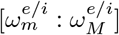:

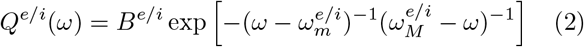

where *B*^*e/i*^ is a normalization factor [31]. *G* denotes the overall coupling constant: this is the main parameter we are going to tune to explore the emergence of different regimes.

The function *Z*(*ϕ*) represents the PRC of each oscillator to a single external pulse. We consider the following PRC for both families of neurons

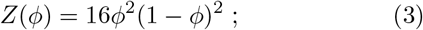

this kind of PRC can be classified as type I [32], and it can be put in correspondence with the dynamical behaviour of an excitable membrane of type I according to the Hodgkin classification [33].

Finally, *C*^*e/i*^ denotes the aggregate input current stimulating excitatory/inhibitory neurons. Under the assumption of a mean-field coupling, *C*^*e/i*^ does not depend on the neuron label,

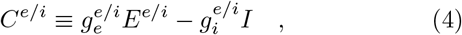

where the 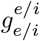 coefficients quantify the specific intra and inter (synaptic) coupling strengths of excitatory and inhibitory populations (here, we have set 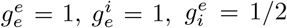 and 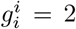). Furthermore, in Eq. (4) *E*^*e/I*^ (*I*) represents the incoming (effective) excitatory (inhibitory) fields, due to the previously delivered excitatory (inhibitory) pulses, namely

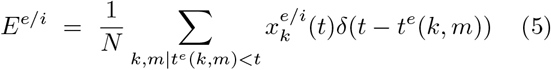

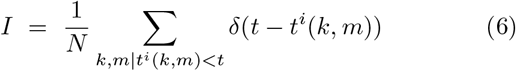

where *t*^*e/i*^(*k, m*) denotes the delivery time of the *m*-th spike by the *k*-th excitatory/inhibitory neuron. 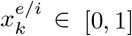 represents the synaptic efficacy of the *k*-th excitatory neuron. If the receiving neuron is inhibitory, 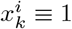, while excitatory-to-excitatory connections are characterized by an adaptive efficacy 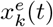. In particular, we assume that its evolution is controlled by an STD mechanism, often invoked in neural systems to simulate the regulation of excitatory activity [25]. According to [24], 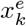 follows the equation

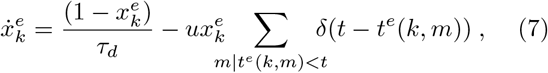

where *t*^*e*^(*k, m*) identifies the emission time of the *m*-th spike by the *k*-th neuron itself. Whenever the neuron spikes, its synaptic efficacy 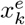 is reduced by a factor (1 − *u*), representing the fraction of resources consumed to produce a post-synaptic spike. So long as the *k*-th excitatory neuron does not spike, the variable 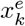 increases towards 1 over a time scale *τ*_*d*_. Altogether, there are 3*N* variables: 2*N* phases characterising the internal state of all neurons, plus *N* variables to quantify the instantaneous value of the excitatory-excitatory synaptic efficacy.

## Numerical Results

In this section, we provide a characterization of the network dynamics for the later interpretation. We start by illustrating the collective behavior. In Fig. S1(a), we plot the time averages of the three fields *I, E*^*e/i*^ vs. the coupling strength *G* for different network sizes from *N* = 8000 to *N* = 32000. The fields appear to be asymptotic (essentially independent of *N*) and stay finite for increasing *G* (they even show a slow decrease): a behavior suggestive of a balanced regime. At *G* = *G*_*θ*_ ≈ 13.5, a dynamical phase transition occurs in the form of a Hopf bifurcation, which separates an AR characterized by constant fields (below *G*_*θ*_), from a regime characterized by oscillating fields (above *G*_*θ*_): the field standard deviations *σ*_*e/i*_ are reported in the inset of Fig. S1(a).

Direct evidence of the oscillatory regime can be appreciated in Fig. S1(b) for *G* = 50, where large irregular peaks are clearly visible in the evolution of the fields, suggesting the presence of a strong synchronization of both excitatory and inhibitory neurons. A complementary view is finally presented in panel (c), where the Fourier power spectra of the excitatory field (those of the inhibitory fields are similar) are plotted for different network sizes. There is no evident dependence on *N*, meaning that the irregularity testified by the broadband structure of the spectra is also an asymptotic phenomenon.

We now shift the focus to the single-neuron level. Fig. S2(a) displays the firing rates of the two populations for *G* = 50: in both cases a plateau is visible, which reveals a mutual synchrony of given groups of neurons at the self-determined frequency *ν*^*e/i*^ ≈ 0.8 (*N* = 16000); a second plateau at its second harmonic is also exhibited by the excitatory neurons.

So far, this phenomenology is very similar to that one reported in [17, 18], where a heterogeneous network of exclusively inhibitory neurons was investigated. However, the single-neuron behavior is, here, significantly more irregular. The coefficients of variation (*CV* s) measuring the level of irregularity in the spiking activity of each neuron [35] instead of being at most around 0.1 as in [17, 18], here they are much larger and closer to the values measured in the brain cortex [36], where the CV is around 1 (see Fig. S2(b)). This suggests and confirms that the simultaneous presence of excitation and inhibition is a crucial ingredient to obtain an irregular microscopic dynamics.

Once more, we wish to stress that the mean-field nature of the model implies that each (excitatory or inhibitory) neuron sees the same field, so that the fluctuations are genuine collective properties rather than statistical fluctuations due to the “sampling” process in a sparse random network.

In order to test the robustness of this scenario, we have simulated also an *annealed* variant of the *quenched* model, where the bare frequencies instead of being fixed forever, are randomly reset (according to the same distribution) every *N* spikes. As shown in Fig. S1 in [34], there are only minor quantitative variations such as the bifurcation point which now occurs for *G*_*θ*_ ≈ 10.5. At the microscopic level, in the annealed model, the neurons are necessarily characterized by the same firing rate and the same *CV* : *ν*^*e*^ = 1.44 ± 0.02 and ⟨*CV* ^*e*^⟩ ≃ 0.43 for the excitatory neurons; *ν*^*i*^ = 0.85 ± 0.03 and ⟨*CV* ^*i*^ ⟩ ≃ 0.84 for the inhibitory ones. Interestingly, the *CV* ‘s are again large, showing that the same scenario emerges in a case (annealed setup) potentially more amenable to an analytic treatment.

## The Homeostatic Mechanism

We now address the question of how such a regime self-sustains, especially for large *G* values. We start by comparing the numerical results with the strictly constant AR, which can be determined from an exact mean field analysis. The details of the calculations in the thermodynamic limit (*N*→∞) are illustrated in the supplemental material [34]; here we simply sketch the procedure and present the final results. Given two tentative values of the (constant) aggregate currents *C*^*e/i*^, one can determine the (constant) single-neuron firing rates by integrating the equations for the phase and the synptic efficacy (see Eqs. (1,7)). Next, one can derive new *C*^*e/i*^-estimates from the firing rates, using Eq. (4): they must be consistent with the initial assumptions. In practice, the analysis amounts to searching a fixed point in the plane (*C*^*e*^, *C*^*i*^), and, once found, to determine the corresponding fields *E*^*e/i*^, *I* from the aggregate currents. The self-consistent results for the fields *E*^*e/i*^, *I* are presented in Figs. S1(a), (see the magenta dashed lines), where we see that they reproduce the numerical observations below the threshold *G*_*θ*_, in agreement with the expectations, since we knew that this regime is asynchronous. In the large *G* limit, the AR still exists and the associated fields stay finite, indicating that this regime is balanced. However, above *G*_*θ*_, the AR is no longer stable. The CC revealed by numerical simulations is also balanced, as visible in Fig. S1 (a), but it is conceptually very different, not only because of the strong fluctuations of fields. This can be appreciated in Fig. S2(c), where the scaled aggregate currents *GC*^*e/i*^ are plotted versus the coupling *G* itself: they remain bounded in the AR (magenta dashed lines), while they diverge in the presence of CC (solid symbols). The asymptotic behavior of the AR can be explained as follows: the finitenes of *GC*^*e/i*^ is a consequence of the finiteness of the fields, combined with an asymmetry between the effective fields of order 1*/G*. This latter balance condition [37] can be satisfied at all because of the presence of synaptic depression, which makes *E*^*e*^ smaller than *E*^*i*^. In the absence of STD, the balanced AR could not exist, except for the special case 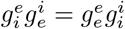.

In the CC regime, the divergence of *GC*^*e/i*^ visible in Fig. S2(c), seemingly contradicts the finiteness of the fields seen in Fig. S1 (a). Consistency is restored by focusing on the role of the PRC. Typically, the efficacy of the incoming spikes, quantified by the PRC, is of order 1 irrespective of the coupling strength. Here, the story is different as shown in Fig. S3(a), where we plot the time and ensemble average 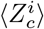 of the PRC, conditioned to the times of the spike arrivals (these are the only instants, where the PRC plays an active role in the neural dynamics).

There, we see that 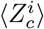 decreases as 1*/G* for large *G* (see black circles) thereby compensating the effect of the increasing coupling strength (in the presence of finite currents). This is an indirect, though clear, indication of an underlying synchronization, since *Z*(*ϕ*) is very small only in the vicinity of the threshold and the reset. Given the phase-like nature of the neuronal variable, one can quantify the degree of synchronization in terms of the modulus of the complex Kuramoto order parameter [2] for the two populations, defined as 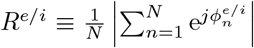 where *j* denotes the imaginary unit. In Fig. S3(a) we show the time averages 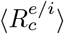 of the order parameters, conditioned to spike emissions, for the two neural populations. In both cases, the parameter approaches 1 with a rate 1*/G*, suggesting a convergence towards perfect synchronization when *G* → + ∞. However, this is not the whole story. In fact, the two unconditioned time averages *R*^*e/i*^ plotted in Fig. S3(b) remain strictly smaller than 1. These two observations thus reveal that the *G*→ ∞ limit is very peculiar. The probability distribution densities (PDFs) of the *R*^*i*^ values, plotted in Fig. S3(c), reveal that a power-law singularity emerges at *R*^*i*^ = 1 for suffi-ciently large *G*. Since the exponent approximately equal to 0.76, is smaller than 1, the singularity is integrable and does not even contribute to the average value of *R*^*i*^, which remains consistently smaller than 1.

Finally, we return to the fluctuations of the spiking activity. Whenever the underlying regime is asynchronous, a large CV means that the neurons operate on average below threshold. The membrane potential (or, equivalently, the phase) is confined to a valley and cannot reach the threshold: this can happen only when a strong fluctuation allows overcoming or removing the barrier. In the present case, the scenario is different: from Fig. S2(c), we see that *GC*^*e*^ | (*GC*^*i*^) | becomes increasingly large and positive (negative) when *G* is increased, suggesting that an excitatory (inhibitory) neuron should find itself much above (below) threshold. However this is not the case, since the large synaptic current values *G* |*C*^*e/i*^| typically contribute to the neuron dynamics at the spike arrivals, when the PRC is small, so that these terms do not sensibly affect the neuron activity.

## Conclusions

We have shown that a mean-field model of pulse-coupled (excitatory and inhibitory) oscillators may give rise a strongly irregular microscopic and macroscopic regime. This is at variance with typical balanced states, where the sparseness contributes to a stochastic-like dynamics by enhancing statistical fluctuations; it is also different from the collective chaos exhibited by chaotic units, as phase-oscillators are never chaotic, even under an external forcing. We claim that the underlying mechanism originates from the selection of a suitable (type-I) PRC shape, which induces a strong but imperfect synchronization. On the one hand, the synchronization stabilizes a balanced regime via a homeostatic mechanism which desentisizes neurons when subject to strong spike bursts. On the other hand, the PRC shape contributes to amplity the tiny differences due to the heterogeneous distribution of bare frequencies, somehow mimicking the sensitivity of standard chaos. The mean-field nature of the model, plus the evidence that the same scenario is observed in an annealed version of the setup (as shown in [34]), together suggest the possibility to develop a theory able to justify our numerical observations. This is the main goal of future work.

AT has received partial support by CY Generations (Grant No ANR-21-EXES-0008) and by the Labex MME-DII (Grant No ANR-11-LBX-0023-01) all part of the French programme “Investissements d’Avenir”. The work has been mainly realized at the Max Planck Institute for the Physics of Complex Systems (Dresden, Germany) as part of the activity of the Advanced Study Group 2016/17 “From Microscopic to Collective Dynamics in Neural Circuits”.

## SUPPLEMENTAL MATERIAL

### A. Asynchronous regime

In the asynchronous regime and for *N*→ ∞, the neurons feel constant fields. It is convenient to write their evolution equation as

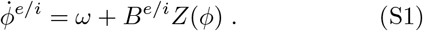

where *B*^*e/i*^ = *GC*^*e/i*^ is defined as

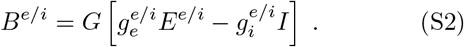

By integrating Eq. (S1), one can determine the interspike interval (ISI)

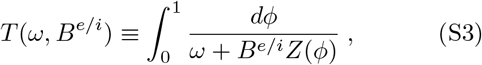

From the ISI, one can obtain the three fields. *E*^*i*^ and *I* are obtained by averaging the inverse of the ISI over the corresponding distributions

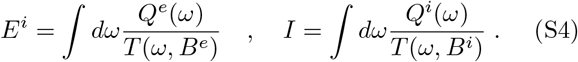

The determination of *E*^*e*^ requires the knowledge of the the synaptic efficacy *x*^*e*^(*ω*) at the spike-emission time,

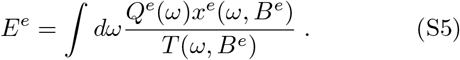

The time-dependent efficacy *x*^*e*^(*t*) obeys the evolution equation

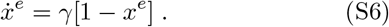

Its general solution is

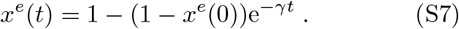

The initial condition *x*^*e*^(0) can be determined self-consistently, by imposing *x*^*e*^(0) = (1 − *u*)*x*^*e*^(*T*^*e*^), where *T*^*e*^ = *T* (*ω, B*^*e*^), so that

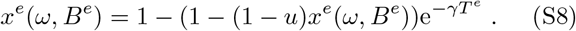

whose explicit solution is

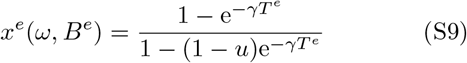

By inserting this expression into Eq. (S5), one can finally determine the last field *E*^*e*^.

Altogether, given *B*^*i*^ and *B*^*e*^, one can determine *E*^*e*^, *E*^*i*^ and *I*. Therefore, we can interpret Eq. (S2) as two self-consistent conditions for two unknowns. In other words, the problem of determining the asyn-chronous state amounts to finding a fixed point in a twodimensional space.

#### 1. Numerics

If we assume that the PRC is *Z*_2_(*x*) = 16*x*^2^(1 − *x*)^2^, the interspike interval Eq. (S3) can be written as (here we drop the superscript for the sake of simplicity)

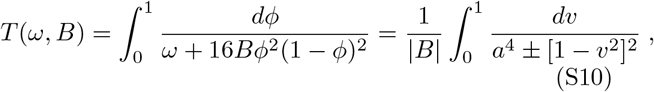

where we have introduced the new variable *v* = 2*ϕ*− 1, while *a*^4^ = *ω/* |*B*|, and the sign in the denominator is the sign of *B*.

The integral can be solved analytically. If the sign is negative, the result is

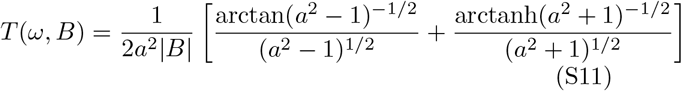

Notice that *a <* 1 means that the slowing down of the coupling is so strong as to block the neuron from firing. Hence, the corresponding neuron does not contribute to the firing rate.

If the sign is positive, the result is

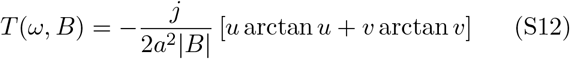

where

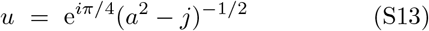

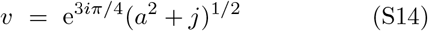

and *j* is the imaginary unit.

#### 2. Large coupling limit

In the large *G* limit, a bounded solution can be obtained if the prefactor of *G* in Eq. (S2) vanishes. The resulting equations imply that, at leading order,

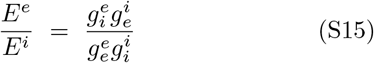

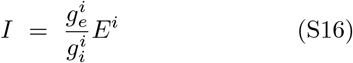

As from Eqs. (S4,S5), both *E*^*e*^ and *E*^*i*^ are function only of *B*^*e*^, which can thus be determined by imposing the condition (S15). Once *B*^*e*^ is estimated, one can determine *B*^*i*^ from Eqs. (S16,S4).

For the parameters employed in the Letter and *u* = 0.50 and *γ* = 0.35 we have found *B*^+^ = 3.628574 and *B*^−^ ≃ 0.619201 and *E*^−^ ≃ 2.256595 and *I* ≃ 1.128311.

### B. Annealed Disorder

In this Section we report the results for the annealed case. The natural frequencies {*ω*^*e/i*^} are still selected from the distribution reported in Eq. (2) of the Letter, but their values, are randomly updated every *N* spikes of the whole network.

**FIG. S1.**
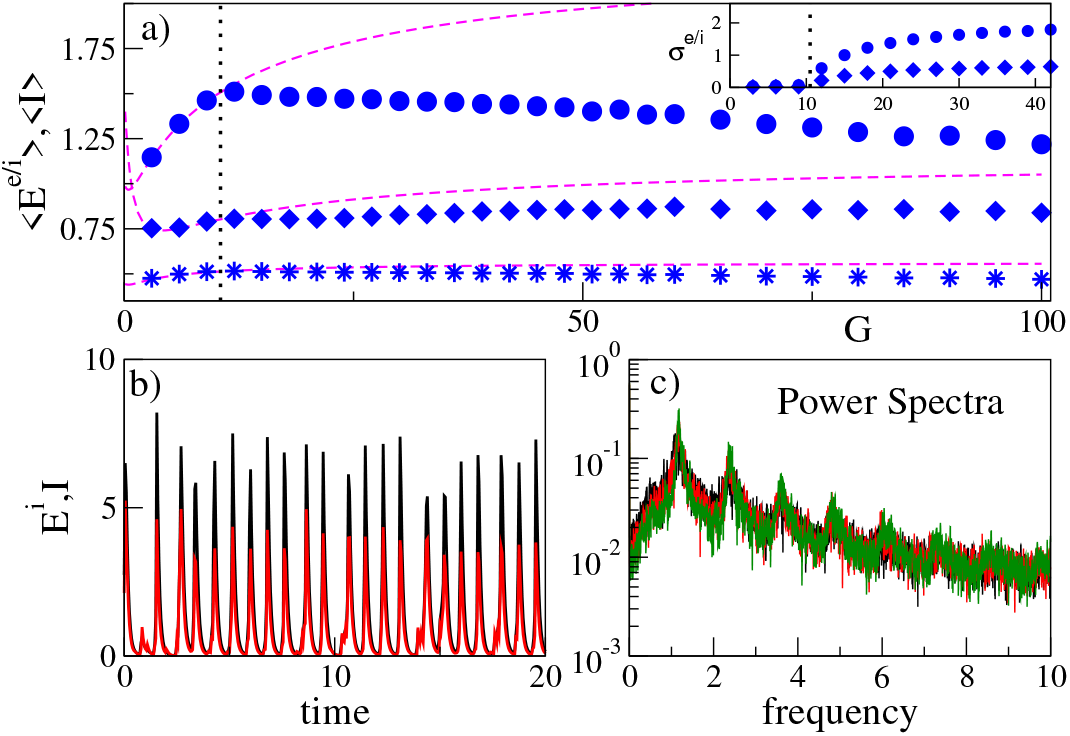
(a) Time averaged synaptic excitatory ⟨*E*^*e/i*^⟩ and inhibitory fields *I* versus *G* for *N* = 16000. The circles denote *E*^*i*^, the diamond ⟨*I* ⟩ and the asterisks *E*^*e*^, the dashed (magenta) lines refer to the exact mean field results for the asynchronous regime. In the inset are reported the standard deviation of the *E*^*i*^ (circles) and *I* fields (diamonds). (b) Excitatory *E*^*i*^ (black) and inhibitory *I* (red) fields versus time. (c) Power spectra for the excitatory *E*^*i*^ field for different system sizes: *N* = 8000 (black), *N* = 16000 (red) and *N* = 32000 (green). Parameters *G* = 50 the spectra refer to an integration time *t* = 5000, after discarding a transient of duration 500. The fields in (b-c) are exponentially filtered with *α* = 10. All the data refer to the annealed case.

In particular, in Fig. S1(a), we plot the time averages of the three fields *I, E*^*e/i*^ versus the coupling strength *G* for different network sizes from *N* = 8000 to *N* = 32000. The field values appear to be essentially independent of *N* and remain finite for increasing *G*: a behavior consistent with a balanced regime. At *G* = *G*_*θ*_ ≈ 10.5, we observe a Hopf bifurcation separating an AR characterized by constant fields (below *G*_*θ*_), from a regime characterized by oscillating fields (above *G*_*θ*_). As clearly visible from the field standard deviations *σ*_*e/i*_ reported in the inset of Fig. S1(a).

Direct evidence of the oscillatory regime can be appreciated in Fig. S1(b) for *G* = 50, where large irregular peaks are clearly visible in the evolution of the fields, suggesting the presence of a strong synchronization of both excitatory and inhibitory neurons. A complementary view is finally presented in panel (c), where the Fourier power spectra of the excitatory field are reported for different network sizes. There is no evident dependence on *N*, meaning that the irregularity testified by the broadband structure of the spectra is also an asymptotic phenomenon.

The self-consistent results for the fields *E*^*e/i*^, *I* in the AR are presented in Figs. S1(a), (magenta dashed lines), where we see that they reproduce the numerical observations below the threshold *G*_*θ*_, in agreement with the expectations. In the large *G* limit, the AR still exists and the associated fields stay finite, indicating that this regime is balanced. However, above *G*_*θ*_, the AR becomes unstable. The CC revealed by numerical simulations is also balanced, as visible in Fig. S1 (a), but it is conceptually very different, not only because of the strong fluctuations of the fields. This can be appreciated in Fig. S2(ab), where the scaled aggregate currents *GC*^*e/i*^ are plotted versus the coupling *G* itself: they remain bounded in the AR (magenta dashed lines), while they diverge in the presence of CC (solid symbols).

**FIG. S2.**
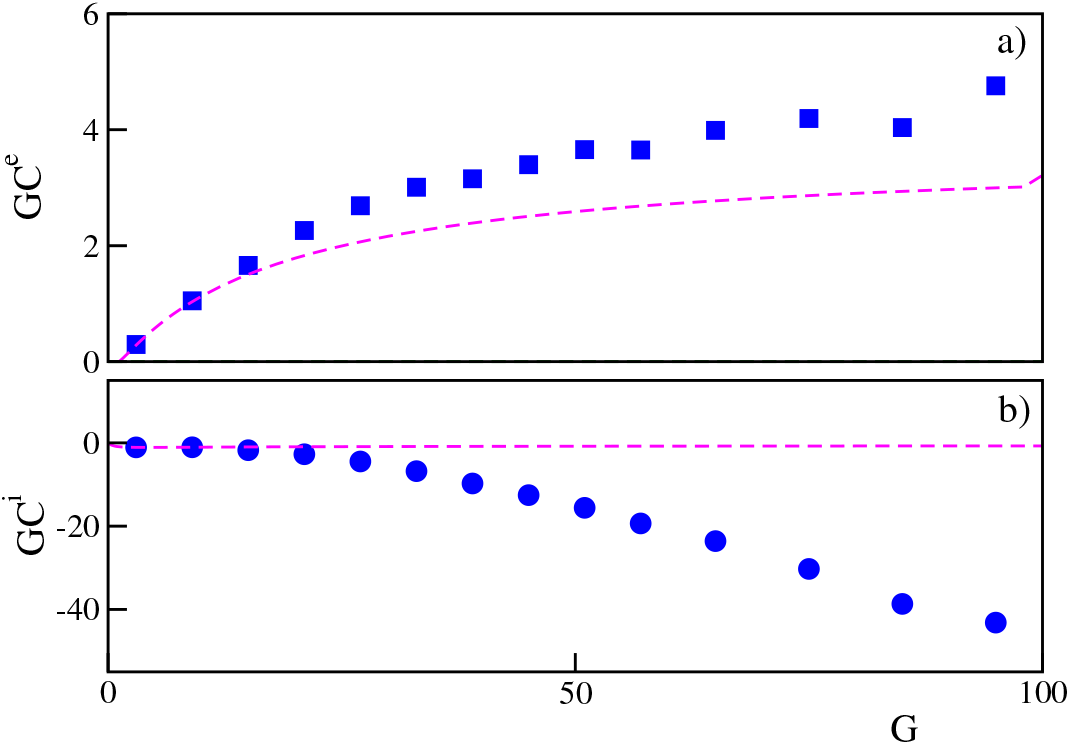
Aggregate synaptic currents *C*^*e/i*^ multiplied by the coupling versus the synaptic coupling itself *G* for excitatory (a) and inhibitory (b) neurons for *N* = 16000. The magenta dashed curves correspond to the exact calculations for the asynchronous regime. The parameters are as in Fig. S1. All the data refer to the annealed case.

In the CC regime, the divergence of *GC*^*e/i*^ visible in Fig. S2(a-b), seemingly contradicts the finiteness of the fields themselves seen in Fig. S1 (a). The consistency can be restored by focusing on the role of the PRC. Typically, the efficacy of the incoming spikes, quantified by the PRC, is of order 1 irrespective of the coupling strength. As shown in Fig. S3(a), where we plot the time and ensemble average 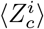 of the PRC, conditioned to the times of the spike arrivals. There, we see that 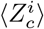 decreases as 1*/G* for large *G* (see black circles) thereby compensating the effect of the increasing coupling strength (in the presence of finite currents). This is an indirect, though clear, indication of an underlying synchronization, since *Z*(*ϕ*) is very small only in the vicinity of the threshold and the reset.

In Fig. S3(a) we show also the time averages 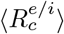 of the order parameters, conditioned to spike emissions, for the two neural populations. In both cases, the parameter approaches one with a rate 1*/G*, suggesting a convergence towards perfect synchronization when *G* → + ∞. However, this is not the whole story. In fact, the two unconditioned time averages ⟨*R*^*e/i*^⟩ plotted in Fig. S3(b) remain strictly smaller than 1.

**FIG. S3.**
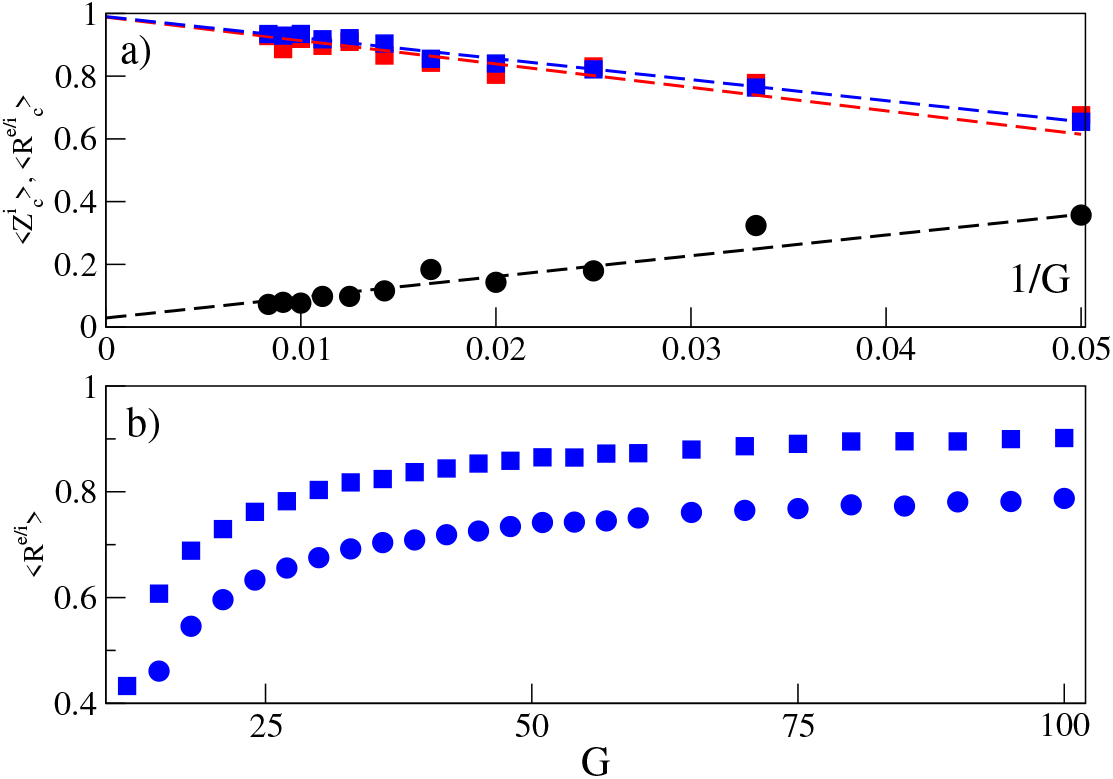
(a) PRC and order parameters at EPSP arrival : 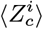 (black circles), 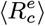 (red squares) and 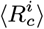 (blue stars) versus 1*/G* for Z2 PRC. We measure the average PRC of a typical inhibitory neuron whenever a EPSP reaches it and at the same time we measure *R*^*e*^ and *R*^*i*^. The dashed lines denote fiting to the data: namely, the black line corresponds to 0.02888 + 6.6138*/G* for a fit t 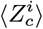 ; the red one to 0.9888 7.4705*/G* for a fit to 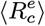 and the blue one to 0.9899 6.703*/G* for a fit to 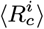. The data refer to *N* = 16000 and to an average over 250000 events. (b) Average value of the Kuramoto order parameter ⟨*R*^*e/i*^⟩ for the excitatory (circles) and inhibitory (squares) population versus *G* for *N* = 16000. Other parameters as in Fig. S1. All the data refer to the annealed case.

Therefore, the overall scenario is not only qualitatively similar to the one observed for the quenched setup, but even quantitatively close.

### C. Network Simulations

The evolution equations Eq. (1) and (7) in the letter are integrated by employing an event driven technique. Since they can be both splitted in an exact evolution between two consecutive speike emissions and and in a nonlinear update of the phases and of the synaptic efficacies at each spike emissione, based on the actual value of the integrated variables as well as for the phases on the values of the the aggregate fields *C*^*e/i*^ and of the PRC 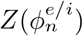. For the evolution of the synaptic efficacies we fix the following parameters *u* = 0.5 and *τ*_*d*_ = 1*/*0.35 = 2.8571 ….

The excitatory and inhibitory fields employed in Fig. 1 (b-c) and in Fig. S1 (b-c) have been obtained by filtering the spike trains with an exponential kernel. In practice, we associate to each spike a an exponential pulse *p*_*α*_(*t*) = *α*e^−αt^ for *t >* 0, and the corresponding excitatory field *E*^*i*^ can be obtained by integrating the following ODE:

**FIG. 1.**
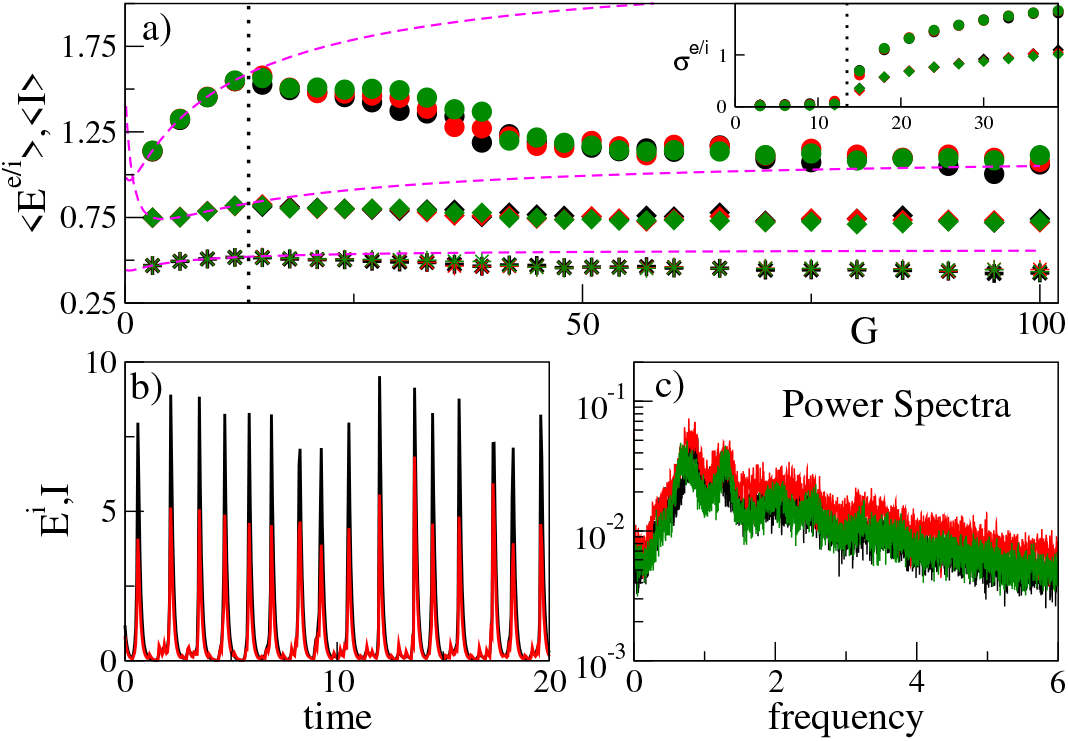
(a) Time averaged synaptic excitatory ⟨*E*^*e/i*^⟩ and inhibitory fields ⟨*I*⟩ versus *G* for different system sizes. The circles denote *E*^*i*^, the diamond *I* and the asterisks *E*^*e*^, the dashed magenta lines refer to the exact results for the AR and the dotted (black) vertical lines denote *G*_*θ*_. In the inset are reported the standard deviations *σ*^*e/i*^ of the *E*^*i*^ (circles) and *I* fields (diamonds). (b) Excitatory *E*^*i*^ (black) and inhibitory *I* (red) fields versus time. (c) Power spectra for the excitatory *E*^*i*^ field for the quenched case for different system sizes. Parameters *G* = 50, the spectra refer to an integration time *t* = 5000, after discarding a transient of duration 500. The fields in (b-c) are filtered with an exponential kernel for more details see [34]. All the data refer to quenched cases. The considered sizes in (a) and (c) are denoted by different colors: namely, *N* = 8000 (black), *N* = 16000 (red) and *N* = 32000 (green).

**FIG. 2.**
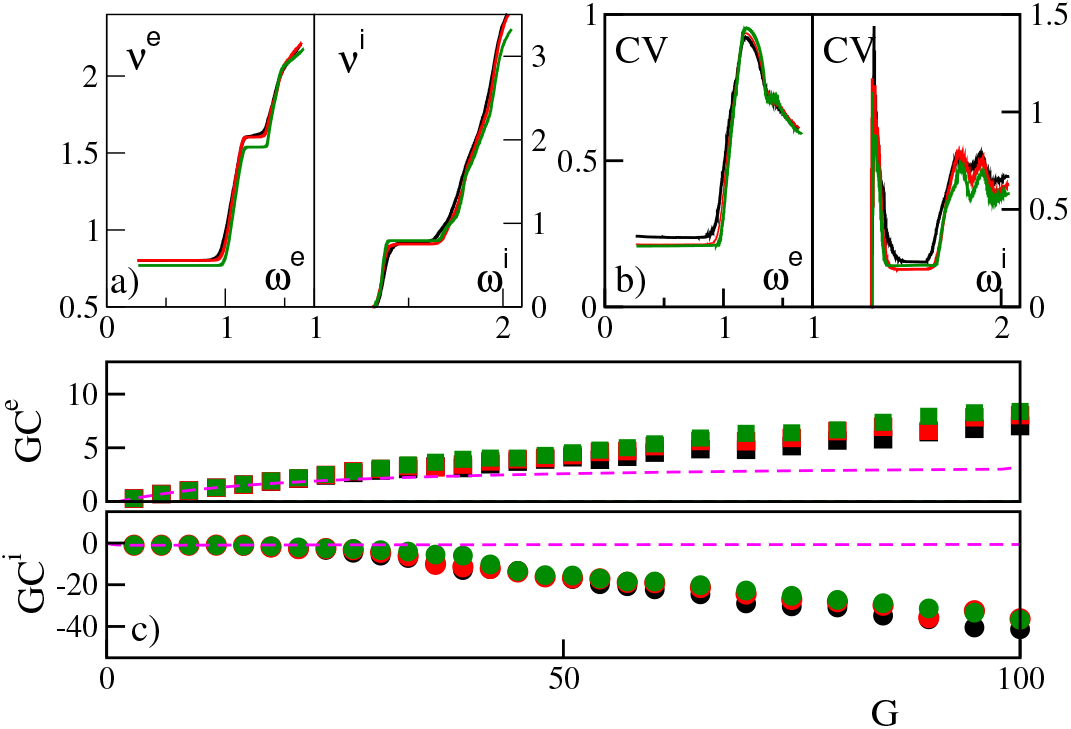
Average firing rates (a) and CV (b) versus their natural frequencies *ω*^*e/i*^ for excitatory (left panels) and inhibitory (right panels) neurons for different system sizes. The data refer to *G* = 50. (c) Aggregate synaptic currents *C*^*e*^ (*C*^*i*^) multiplied by the coupling versus the synaptic coupling itself *G* for excitatory (inhibitory) neurons for different system sizes are shown in the upper (lower) panel. The magenta dashed curves correspond to the exact calculations for the AR. All the data refer to quenched cases. The considered sizes are denoted by the same colors as in Fig. S1 (a).

**FIG. 3.**
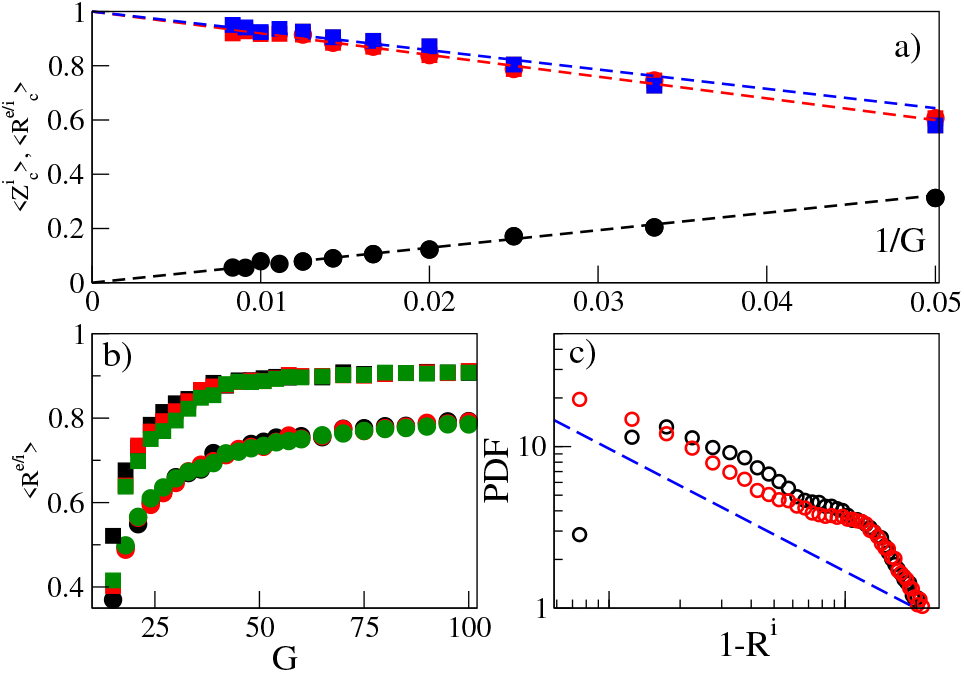
(a) Conditional PRC and order parameters: *Z*^*i*^ (black circles) 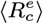 (red squares) and 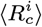 (blue squares) versus 1*/G*. We measure the average PRC of a typical inhibitory neuron whenever an excitatory pulse reaches it and at the same time we measure *R*^*e*^ and *R*^*i*^. The data refer to *N* = 16000 and to an average over 250000 events. (b) Average value of the Kuramoto order parameter for the excitatory (circles) and inhibitory (squares) population versus *G* for various systems sizes: the sizes are encoded as in Fig. S1 (a). (c) PDF of 1 *R*^*i*^ for *G* = 50 (black circles) and *G* = 100 (red circles), the dashed blue line denotes a power-law decay with exponent 0.76. The data correspond to *N* = 16000. Parameters *K* = 0.5.

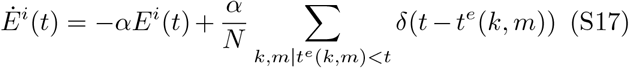

where *t*^*e*^(*k, m*) denotes the delivery time of the *m*-th spike by the *k*-th excitory neuron. Analogously we have obtained the filtered inhibitory fields. We have always set *α* = 10.

## References

[1] A. T. Winfree, The Geometry of Biological Time, vol. 12 of Interdisciplinary Applied Mathematics (Springer-Verlag New York, 2001), 2nd ed.

[2] Y. Kuramoto, Chemical oscillations, waves, and turbulence, vol. 19 (Springer Science & Business Media, 2012).

[3] R. E. Mirollo and S. H. Strogatz, SIAM Journal on Applied Mathematics 50, 1645 (1990).

[4] C. Liu, D. R. Weaver, S. H. Strogatz, and S. M. Reppert, Cell 91, 855 (1997).

[5] H. Haken, Brain dynamics: synchronization and activity patterns in pulse-coupled neural nets with delays and noise (Springer Science & Business Media, 2006).

[6] F. Dörfler and F. Bullo, Automatica 50, 1539 (2014).

[7] A. Pikovsky, M. Rosenblum, J. Kurths, and A. Synchronization, Self 2, 3 (2001).

[8] S. H. Strogatz, Sync: How order emerges from chaos in the universe, nature, and daily life (Hachette UK, 2012).

[9] P. C. Matthews and S. H. Strogatz, Physical review letters 65, 1701 (1990).

[10] K. Kaneko, Physical review letters 65, 1391 (1990).

[11] V. Hakim and W.-J. Rappel, Physical Review A 46, R7347 (1992).

[12] N. Nakagawa and Y. Kuramoto, Progress of Theoretical Physics 89, 313 (1993).

[13] S. Olmi, A. Politi, and A. Torcini, Europhysics Letters 92, 60007 (2011).

[14] D. Pazó and E. Montbrió, Physical Review X 4, 011009 (2014).

[15] D. M. Abrams and S. H. Strogatz, Physical review letters 93, 174102 (2004).

[16] D. Pazó and E. Montbrió, Physical review letters 116, 238101 (2016).

[17] S. Luccioli and A. Politi, Phys. Rev. Lett. 105, 158104 (2010).

[18] E. Ullner and A. Politi, Physical Review X 6, 011015 (2016).

[19] D. Sherrington and S. Kirkpatrick, Physical review letters 35, 1792 (1975).

[20] H. Daido, Physical review letters 68, 1073 (1992).

[21] C. van Vreeswijk and H. Sompolinsky, Science 274, 1724 (1996).

[22] A. Politi and A. Torcini, Chaos: An Interdisciplinary Journal of Nonlinear Science 34 (2024).

[23] W. R. Softky and C. Koch (1992).

[24] M. Tsodyks, K. Pawelzik, and H. Markram, Neural computation 10, 821 (1998).

[25] M. Tsodyks and S. Wu, Scholarpedia 8, 3153 (2013).

[26] G. Mongillo, D. Hansel, and C. Van Vreeswijk, Physical review letters 108, 158101 (2012).

[27] L. F. Abbott and C. Van Vreeswijk, Physical Review E 48, 1483 (1993).

[28] M. Timme, F. Wolf, and T. Geisel, Physical review letters 89, 258701 (2002).

[29] M. Denker, M. Timme, M. Diesmann, F. Wolf, and T. Geisel, Physical review letters 92, 074103 (2004).

[30] C. van Vreeswijk, Physical Review E 54, 5522 (1996).

[31] Here, we have set 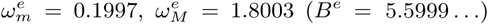, and 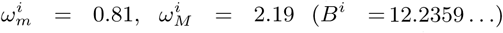. The two choices correspond to an average 1 and 1.5, respectively and have the same standard deviation as flat distributions of total width 1 and 0.8, respectively.

[32] D. Hansel, G. Mato, and C. Meunier, Neural computation 7, 307 (1995).

[33] C. C. Canavier, Scholarpedia 1, 1332 (2006).

[34] See Supplemental Material at [URL will be inserted by publisher] for details on the self-consistent analysis for the asynchronous regime, for the results on the annealed case and details on the performed simulations.

[35] The coefficient of variation CV of a certain neuron is the ratio between the standard deviation and the mean of the inter-spike intervals associated to the train of spikes emitted by the considered neuron. CV = 0 corresponds to a periodic activity, while CV = 1 is observable for a Poissonian spike train.

[36] W. R. Softky and C. Koch, Journal of neuroscience 13, 334 (1993).

[37] Usually in the dynamical balance theory [21] the coupling strenght scales as 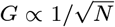 therefore the difference of the effective fields, contained in Ce/i, grows as 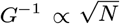 to obtain the balancing.

